# Imbalance in the response of pre- and post-synaptic components to amyloidopathy

**DOI:** 10.1101/599399

**Authors:** Terri-Leigh Stephen, Francesco Tamagnini, Judith Piegsa, Katherine Sung, Joshua Harvey, Alice Oliver-Evans, Tracey K. Murray, Zeshan Ahmed, Michael L. Hutton, Andrew Randall, Michael J. O’Neill, Johanna S. Jackson

**Author notes:** co-first author (Terri-Leigh Stephen, Francesco Tamagnini).

## Abstract

Alzheimer’s disease (AD)-associated synaptic dysfunction drives the progression of pathology from its earliest stages. Aβ species, both soluble and in plaque deposits, have been causally related to the progressive, structural and functional impairments observed in AD. It is, however, still unclear how Aβ plaques develop over time and how they progressively affect local synapse density and turnover. Here we observed, in a mouse model of AD, that Aβ plaques grow faster in the earlier stages of the disease and if their initial area is > 500 µm^2^; this may be due to deposition occurring in the diffuse part of the plaque. In addition, synaptic turnover is higher in the presence of amyloid pathology and this is paralleled by a reduction in pre-but not post-synaptic densities. Plaque proximity does not appear to have an impact on synaptic dynamics. These observations indicate an imbalance in the response of the pre- and post-synaptic terminals and that therapeutics, alongside targeting the underlying pathology, need to address changes in synapse dynamics.

## Introduction

Alzheimer’s disease (AD) is the world’s leading cause of dementia and is thought to be, primarily a synaptopathy (Kerrigan and Randall, 2013; Selkoe, 2002). Indeed, synaptic loss and altered connectivity occurs in the earliest stages of AD, preceding its histopathological hallmarks and possibly driving the progression of cognitive decline (Canuet et al., 2015; Grandjean et al., 2014; Kim et al., 2013). Mutations in the gene encoding the amyloid precursor protein (APP) carry the greatest incidence of early-onset familial AD (FAD), along with presenilin 1 and 2 (Bittner et al., 2012). APP is widely expressed in neurons and is thought to be responsible for synapse formation and repair (Randall et al., 2010). Animal models with human APP mutations exhibit synapse loss and dysfunction, gliosis (Jackson et al., 2017) cognitive impairments (Demattos et al., 2012) and eventual neuron loss [39] alongside the progressive accumulation of pathogenic Aβ species (Hong et al., 2016).

To further understand synapse dynamics in amyloidopathy, we assessed plaque and synapse dynamics in the J20 model, which has human APP mutations consistent with FAD (Mullan et al., 1992; Murrell et al., 1991). In this model, soluble Aβ is apparent at 2 months of age, with the first plaques starting to accumulate in the 5^th^ month, in some animals, and in all animals by the 10^th^ month (Mucke et al., 2000). From the moment Aβ plaques start to appear, pathogenic amyloid species are not uniformly distributed across the brain parenchyma. Consequently, there may be differing effects on synapse structure and function in relation to their proximity to plaques (Spires et al., 2005). Previous observations identified an inverse relationship between dendritic spine density and amyloid plaque pathology [35,21].

As Aβ deposits represent a reservoir of pathogenic amyloid species, we first investigated plaque growth rates in relation to their initial area, plaque region and animal’s age in vivo, using the amyloid-binding dye methoxy-XO4 (Klunk et al., 2002), which labels both condensed cores and the surrounding diffuse fibrillar Aβ subspecies (Wang et al., 2016). In addition, we hypothesized that synaptic density and turnover in the neocortex are altered in amyloidopathy between 7 and 10 months of age and differentially affected by plaque proximity.

## Results

To measure synapse dynamics in relation to plaque deposition in AD-associated amyloidopathy, we used *in vivo* two-photon imaging of methoxy-X-O4-labelled plaques and GFP-expressing pyramidal neurons in the somatosensory cortex (SSC) of J20 mice and WT littermate controls (**Fig. 1a**). The same regions of interest were imaged longitudinally on a weekly basis. We studied two groups of J20 animals at different ages ranging from early (30-42 weeks/7-10 months old) to late stages (49-61 weeks/11-14 months old) (**Fig. 1b**) of the disease. WT littermates were used as a negative control cohort for the younger age-point (30-42 weeks/old) only. Both of the stages evaluated were characterized by the presence of pathogenic Aβ species and synapse loss in transgenic animals (Hong et al., 2016; Mucke et al., 2000).

**Figure 1.**
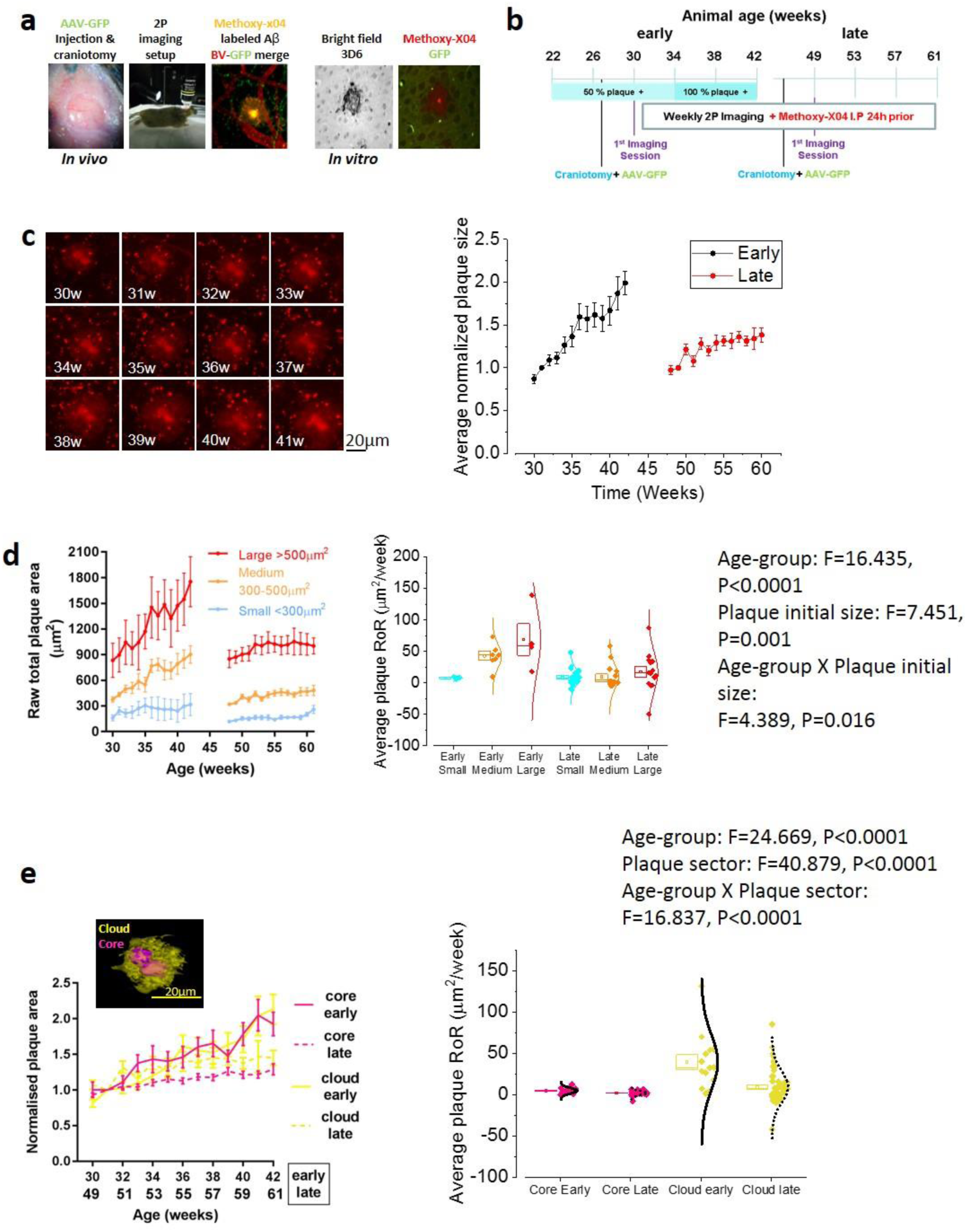
Amyloid plaque dynamics in Alzheimer’s disease - early vs late stage. (**a**) *In vivo* two-photon imaging set-up with example image of adeno-associated virus (AAV) GFP-transduced pyramidal neurons in layer 2/3 of the somatosensory cortex (SSC) (green) alongside blood vessels labelled with Texas-red (red) and plaques labelled with methoxy-X04 (yellow). Right inset images show colocalization of the Aβ antibody 3D6 and methoxy-X04 *in vitro*. (**b**) Experimental timeline of early and late J20 groups. The early group was imaged between 7 and 10 months (30-42 weeks) while the late stage group was imaged between 11 and 14 months of age (49-61 weeks). (**c**) Representative image series showing amyloid plaque growth starting at 30 weeks (left panel) and quantification of total plaque growth (right panel) normalized to the second imaging session (one-way ANOVA, early p = 0.0002 and late p = 0.0002). Plaques grew faster in younger mice (unpaired t-test; p < 0.0001). **(d)** *Left panel:* raw total plaque area (µm^2^) in the early (30-42 weeks) and late (49-61 weeks) groups separated into size categories; small (< 200 µm^2^, blue), medium (200-500 µm^2^, orange) and large (> 500 µm^2^, red). Small plaques (Two-way ANOVA, Factor: Plaque’s initial size, Variable: slope, F=7.451, p = 0.001) in younger animals (Two-way ANOVA, Factor: age, Variable: slope, F=16.435, p < 0.0001) grew faster. *Right panel:* Scatter-box plot showing the plaque-to-plaque growth rate (Rate of Rise: RoR, units: µm^2^/week) grouped per age and plaque’s initial size. Two-way ANOVA revealed that both age and plaque initial size affect the plaque RoR (p = 0.02). Post-hoc Bonferroni corrected t-tests revealed that larger plaques grow faster than small plaques in the early age-group (Two-way ANOVA, Factor: age x Plaque’s initial size, Variable: slope, F=4.389, **P=0.0016**, see SI Table 1 for all p values). In this and all the subsequent scatter-box plot figures, the box represents the mean (small square) the SEM boundaries (top and bottom line) and the median; the distribution line on the right shows the normality of the distribution for each data population. (**e**) *Left panel:* Normalized area of the plaque cloud (yellow) and core (pink) in the early (broken lines) and late groups (solid lines). *Right panel:* scatter plot showing plaque-to-plaque average RoR, grouped per age and plaque region (core or cloud). The cloud grew faster than the core (Bonferroni’s corrected unpaired t-test; **P<0.0001**) and the plaques in younger mice grow faster only at the level of the cloud (Bonferroni’s corrected unpaired t-test; **P<0.0001**). Early group n = 13 plaques, late group n = 58 plaques. Unless otherwise stated, data are presented as means ± SEM.

### Amyloid plaque growth-rate is faster in younger animals and for bigger plaques

The total area of Aβ plaques increased in both early and late cohorts (**Fig. 1c** and **S.Fig. 1; early *p* = 0.0002 and 1b late *p* = 0.0002**). The plaque-to-plaque average growth rate (Rate of Rise: RoR, units: µm^2^/week) was measured as the slope of a linear function fit for each plaque’s surface area in time. We observed that Aβ plaques grew faster in younger animals (**Fig 1c right; *p* <0.0001**). As the plaques were not homogenous in size, we separated them into three groups, based on the initial raw total area: small (<300 µm^2^) medium (300-500 µm^2^) and large (>500 µm^2^). We observed (**Fig 1d**) that plaques which are initially large (***p*=0.001**) and that are in younger animals (***p*<0.0001**) grew faster; in addition, age and plaque initial size interact in affecting plaque growth rate (***p*=0.0016**). For the complete set of pairwise comparisons, see SI-Table 1.

We separately investigated the growth rate of the plaque’s dense core and diffuse cloud, across the two age-points investigated (Fig 1e). We observed that the age-point and the plaque’s sector both affected and interacted with each other in affecting the plaque’s growth rate (**See insert in Fig 1e for F and P values**): the cloud grew faster than the core at both the early (***p*<0.0001**) and late (Bonferroni’s corrected unpaired t-test; ***p*=0.008**) age-points. However, the core’s growth rate was not different between age-points (Bonferroni’s corrected unpaired t-test; ***p*=0.542**), while the cloud’s growth rate was faster in the early group, compared to the late one (***p*<0.0001**). For the complete set of pairwise comparisons, see SI-Table 2. From these results, we can conclude that plaques grow faster in younger animals and if they have a big, diffuse cloud. This is presumably due to amyloid β aggregation occurring quicker at the level of the diffuse plaque zone.

### Axonal terminaux boutons but not dendritic spines are reduced in J20s

Hippocampal synapse loss has been observed in the J20 model as early as 3 months (Hong et al., 2016). However, the subtle differences between pre- and post-synaptic components have not been elucidated in relation to significant levels of amyloid, or in the cortex. Using methods we have previously developed for studying synapse turnover (Jackson et al., 2017), we examined whether the pre- and post-synaptic components (axonal terminaux boutons (TBs) and dendritic spines, respectively) were altered in the J20 model, in an age range exhibiting significant plaque accumulation (between 7 and 10 months/30 and 42 weeks), compared to WT littermates (**Fig. 2a**). There was no overall difference in spine density (**Fig. 2b, *p* = 1**), however, we observed a significant decrease in the density of TBs in J20s compared to WT controls in the SSC (**Fig. 2c, *p* = 0.04**).

**Figure 2.**
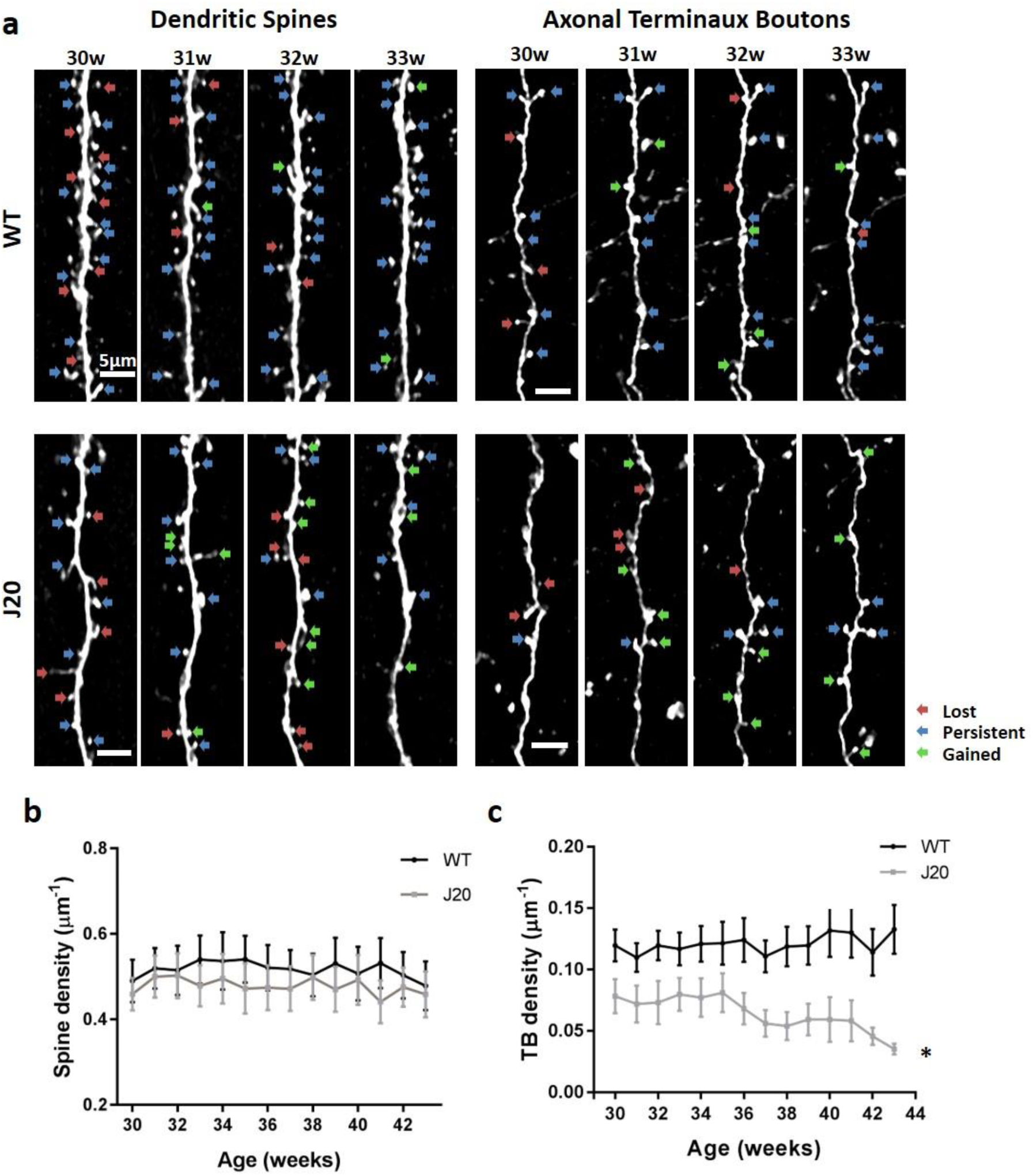
Axonal terminaux boutons but not dendritic spines are reduced in J20s. (**a**) Representative maximum projections of dendritic spines and axonal TBs in J20 animals and littermate controls (WT) at 30-33 weeks of age. Red arrows show lost, blue arrows show persistent and green arrows show gained spines/boutons. (**b**) Quantification of dendritic spine density in J20s and WT control animals showing no significant difference (two-way repeated measure (RM) ANOVA genotype x age *F*_(605, 13)_ = 0.32,(p = 1). (**c)** Quantification of TB density shows a progressive loss with age compared to WT controls (two-way RM ANOVA genotype x age *F*_(517,13)_ = 1.8, p = 0.04). J20 spines n = 21, WT spines n = 34, J20 TBs n = 16, WT boutons n = 36. Error bars represent SEM.

### The stability of dendritic spines and axonal boutons is altered in J20s

Continual addition and loss of synapses is thought to underlie the fine tuning of neuronal function to match cognitive demands [(Grillo et al., 2013; Holtmaat et al., 2006; Majewska et al., 2006; Trachtenberg et al., 2002). Disruption of synapse turnover or stability is thought to occur in disease, potentially indicative of the early stages of dementia (Cruz-Martín et al., 2010; Jackson et al., 2017; Murmu et al., 2013),27]. Thus, to further examine how synapse dynamics are altered in our model of amyloidopathy, spine and TB survival, as a fraction of the first-time point, was initially analyzed. There was a significant decrease in the survival fraction of both spines and TBs compared to WT control animals (**Fig. 3a, spines *p* = 0.04 and TBs *p* < 0.001**). The turnover ratio (TOR) was also disrupted, as there was a significant increase in the TOR for both dendritic spines and TBs in the J20 group (**Fig. 3c, spines *p* = 0.04 and TBs *p* < 0.001**). In addition, comparing the turnover of spines and TBs, as an average of all the time points, indicated a significantly higher turnover of TBs compared to spines in J20s (**Fig. 3e, *p* = 0.01**). On further analysis, this disruption in turnover was driven by an increase in gained spines while the number of lost spines was unaffected (**Fig. 3f, gains *p* = 0.02 losses *p* = 0.06**). The increased turnover of axonal TBs was driven by both gained and lost boutons (**Fig. 3g, gains *p* = 0.02 losses *p* = 0.004**). These observations reveal that, in the J20 model, both spines and TBs are less stable and the balance of turnover is significantly disrupted at the level of both the pre- and post-synapse.

**Figure 3.**
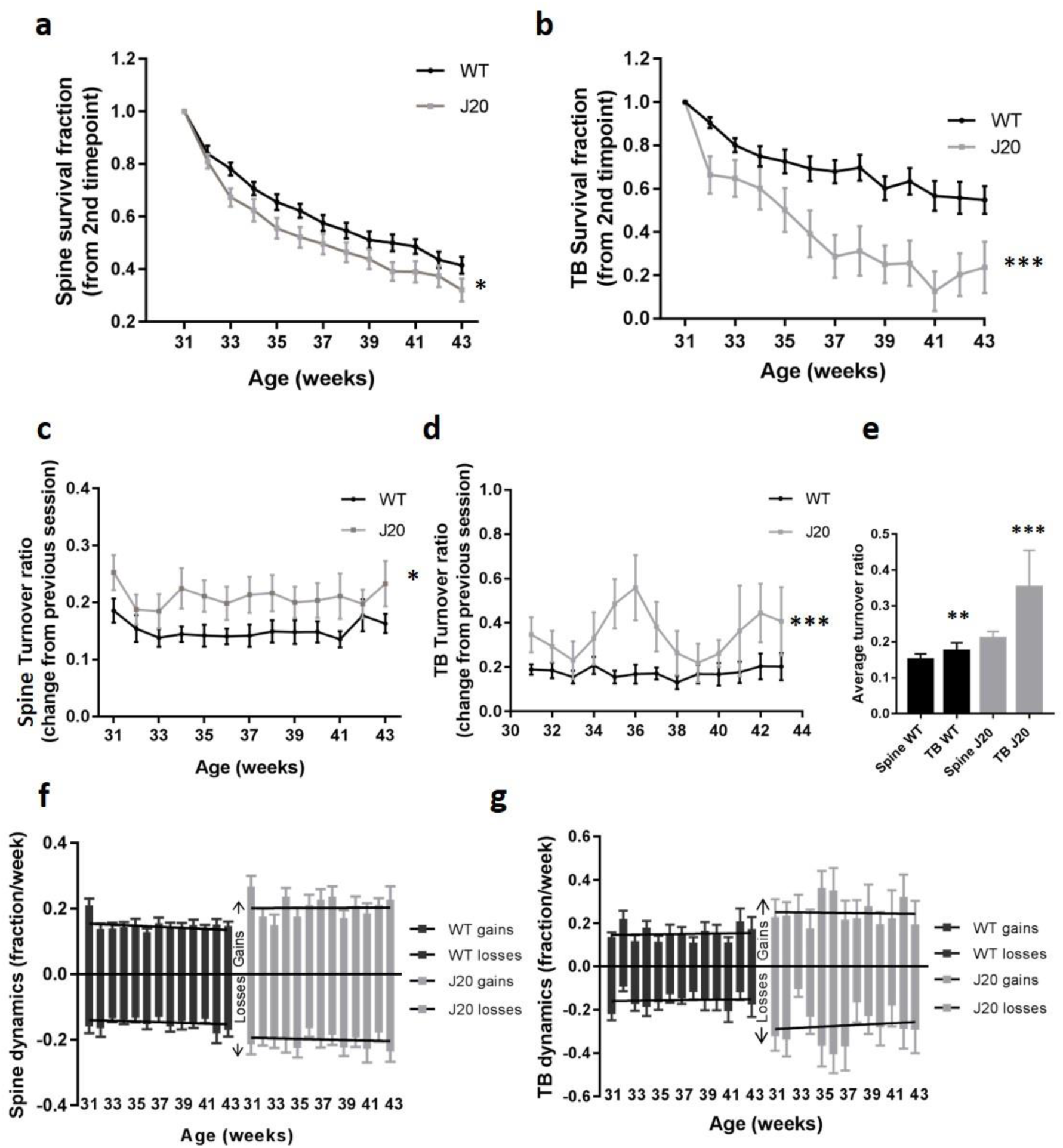
The stability of dendritic spines and axonal terminaux boutons are altered in J20s. (**a-b**) Quantification of spine (**a;** two-way RM ANOVA genotype *F*_(540,1)_ = 4.4, p = 0.04) and TB (**b;** two-way RM ANOVA genotype *F*_(461,1)_ = 13.7, p < 0.001) survival, as a fraction of the first time-point (30 weeks), in J20s and WT controls. (**c-d**) TOR, showing the change compared to the previous session, of spines (**c;** two-way RM ANOVA genotype *F*_(552,1)_ = 4.4, p = 0.04) and TBs (**d;** two-way RM ANOVA genotype *F*_(471,12)_ = 112.3, p < 0.001) in J20s and WT controls. (**e**) Average total turnover across all time points showed a significant difference in spine and TB TOR in the J20 group but not in the WTs (two-way ANOVA *F*_(103,1)_ = 6.3, p = 0.01). (**f-g**) Gains and losses of dendritic spines (**f**; two-way RM ANOVA gains genotype *F*_(552,1)_ = 5.6, p = 0.02; losses genotype *F*_(552,1)_ = 3.8, p = 0.06) and TBs (**g**; two-way RM ANOVA gains genotype *F*_(482,1)_ = 5.8, p = 0.02; losses genotype x age *F*_(482,12)_ = 2.5, p = 0.004) in J20s compared to WT controls. J20 spines n = 21, WT spines n = 34, J20 TBs n = 16, WT TBs n = 36. Error bars represent SEM.

### Alterations in synapse density and stability is not affected by plaque proximity

As previous studies have shown that synapse density is negatively correlated to the proximity of amyloid plaques in other amyloid models (Tg2576 with the Swedish APP mutation) [33], we thus aimed to assess this in the J20 amyloid model. The proximity of the nearest plaque was measured for each axon or dendrite (**Fig. 4a**). Neurites were considered close to a plaque if the nearest plaque was within 300 µm. The densities of both axonal and dendritic synapse components were not differentially affected if the nearest plaque was less than 300 µm away when compared to those without a plaque nearby at any time point (**Fig. 4b, *p* = 0.5 and c, *p* = 0.2**). Notably, new plaques appeared during our longitudinal imaging periods, meaning plaques were at different stages of maturity and of different sizes. Consequently, we decided to consider a neuronal process as close to a newly formed plaque at five weeks post-plaque appearance. However, newly formed plaque distance did not significantly correlate with the spine or TB density (**Fig. 4e, *p* = 0.7 and *p* = 0.2, respectively)**. The stability (**Fig 4e, spine *p* = 0.7 and TB *p* = 0.08)** and turnover (**Fig. 4f, spine *p* = 1 and TB *p* = 0.2**) of spines and TBs was also unaffected by their proximity to the nearest plaque. Post-mortem evaluation of synaptophysin and PSD-95 puncta close to plaques confirmed that, even within 50 µm of plaques (**Fig. 4g**), pre-synaptic terminals (**Fig. 4h, *p* < 0.001)** were lost compared to WTs whereas post-synaptic spines were unaffected (**Fig. 4i, *p* > 0.05**).

**Figure 4.**
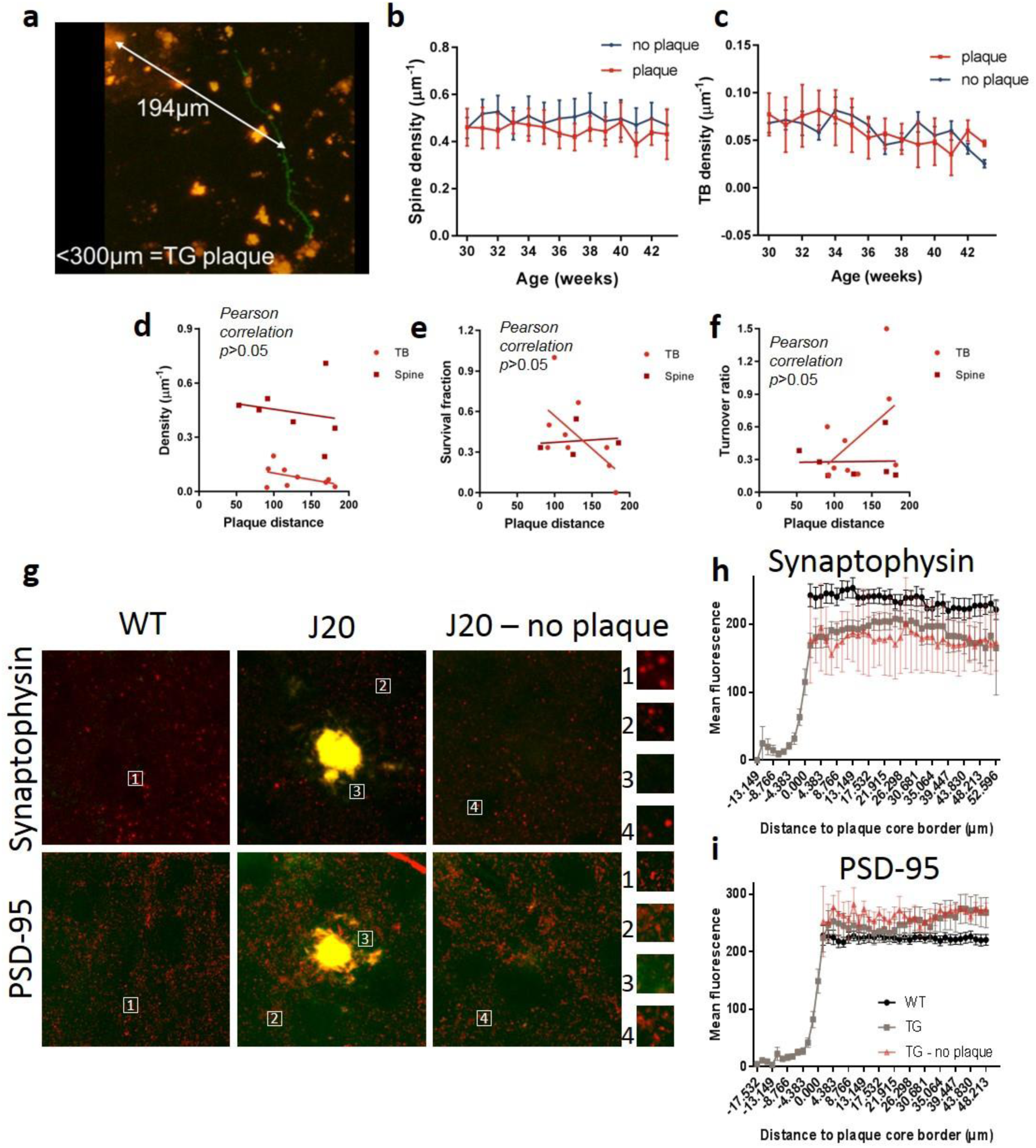
The changes in spine and bouton dynamics are not affected by plaque proximity or maturity. (**a**) The distance of each neurite to the center of the nearest plaque was measured. (**b-c**) Dendrites (**b**; two-way RM ANOVA plaque *F*_(204,1)_ = 0.5, p = 0.5) and axons (**c;** two-way RM ANOVA plaque *F*_(141,11)_ = 1.7, p = 0.2) within 300 µm of a plaque did not have a significant difference in synapse density compared to those further away from a plaque (> 300 µm). (**d-f**) Plaque distances were measured for each neurite at five weeks post-plaque appearance, to control for the appearance of new plaques. This did not affect spine (**d**; Pearson correlation p = 0.7) or TB density (**d;** Pearson correlation p = 0.2), survival fraction (**e;** Pearson correlation, spines p = 0.7, TBs p = 0.08) or turnover ratio (**f;** Pearson correlation, spines p = 1, TBs p = 0.2). (**g**) CLARITY-cleared brain slices were stained for synaptophysin and PSD-95 and the immunoreactivity around plaques quantified. (**h**) Synaptophysin intensity was significantly reduced within 50 µm of plaques with a significant interaction between distance to plaque and group (two-way RM ANOVA *F*_(949,64)_ = 2.5, p < 0.001). (**i**) PSD-95 puncta intensity was not altered close to plaques (two-way RM ANOVA *F*_(1438,64)_ = 0.5, p > 0.05).

## Discussion

The loss of synapses in AD has been observed in humans and several mouse models (Selkoe, 2002; Walsh and Selkoe, 2004), however, the specific, time-dependent dynamics of synaptic alterations warrants further investigation. Further, while the growth of amyloid plaques has been previously characterized *in vivo* (Burgold et al., 2014), the growth dynamics of plaques, as a function of their size and age, and the relationship between plaque growth and synapse dynamics still remains to be determined. Subtle changes in fine plaque and synaptic detail, and their interactions over time, captured here using *in vivo* multiphoton imaging, cannot readily be studied using conventional post-mortem analyses.

In this current study, we present three main findings: firstly, amyloid plaques grow at a faster rate if they are large and in young animals, secondly axonal boutons, but not dendritic spines, are lost in amyloidopathy and thirdly, the stability of both synaptic structures is compromised in the J20 transgenic model. The age- and size-dependent differences in plaque growth rate in J20 mice may be due to deposition dynamics occurring in the diffuse part of the plaque, rather than the dense core. This suggests that the main area of plaque growth may be the diffuse cloud: the slow growth of small plaques may be due to the reduced extension of the cloud. Age, plaque size and extension of the diffuse cloud, may be of importance when determining drug interventions (Demattos et al., 2012).

Whilst we characterized the growth of individual plaques in this study, the overall density of plaques in the SSC also increased over this time period (Mucke et al., 2000). The growth of individual plaques was affected by the initial size, especially in the younger animals, however, the higher density of plaques in the older animals (late group) may account for the reduced growth with age (Burgold et al., 2014). Furthermore, our data suggests that the main area of plaque growth may be the diffuse cloud. Whilst it is possible that the different groups may have differing rates of Aβ production, the aggregated Aβ in both groups would be deposited either as a new plaque or join an existing plaque. In young animals, where plaques are developing, the Aβ would join the relatively few existing plaques, reflected by a higher rate of growth in these plaques, especially in the diffuse plaque cloud. However, in older animals, where there are more plaques, new Aβ would be distributed amongst the many plaques present so each plaque would show a slower, possibly saturated, rate of growth. We hypothesize that the cloud consists of loosely packed fiber-like structures that extend outward from the denser packed core, as described previously (Wang et al., 2016). Recent literature has proposed a link between plaque compaction and microglial-dependent plaque interactions that have a direct effect on the degree of neurite damage (Yuan et al., 2016). Indeed, a study by Condello and collegues has shown that rapidly growing plaques are associated with greater levels of local neuronal dystrophy and that this damage plateaus as plaque growth rate slows (Condello et al., 2011). Of further note, microglia have been shown to form a physical ‘barrier’ around plaques that limits their expansion and result in smaller, more compact plaques (Condello et al., 2015). This could explain the heterogeneity in plaque growth rate, where the growth rate of smaller plaques might be limited by microglia. It should be noted that the older group of animals received a control compound in a drug discovery study which was found to not have any effects on amyloid pathology although other more subtle effects cannot be ruled out.

This is the first study of its kind to quantify the growth of different components of amyloid plaques (specifically comparing the cloud and the dense core) at different time points using this temporal resolution. Overall, these observations help us to understand how amyloid pathology progresses over time and may help to identify patients for newly developed therapies. This could potentially be achieved by the validation of the detection of single Aβ plaques in the retina as a selective and specific diagnostic tool for AD (Jiang et al., 2016). Aβ plaque number, rate of growth (both in number and size), cloud:core surface ratio and the correlation of these parameters with the cognitive state of the patient, may provide vital information on the state of progression of the disorder and what therapeutic strategy may be the most effective, for individual patients.

To date, very few studies have looked at dendritic spine density in the J20 model. A 36% reduction was observed in the hippocampus at 11 months (Moolman et al., 2004), however, equivalent measurements were not made in the cortex. Interestingly, Hong *et al*. reported a significant decrease in hippocampal PSD-95 puncta in J20s, at 3 months of age, without a change in synaptophysin staining in *ex vivo* sections [13]. However, Mucke *et al.*, who originally characterized the J20 model, reported a loss of synaptophysin terminals not correlated with plaque load suggesting that plaques are not the cause of synapse loss, consistent with our findings. In the Tg2576 model, Spires *et al.* showed a decrease in cortical spines (Spires et al., 2005) and Bittner *et al*. showed a decrease in cortical and hippocampal spines in 3xTg mice (both within and > 50 µm from plaques) (Bittner et al., 2012). Here, however, we show that dendritic spine density was unaffected in the cortex (both *in vivo* and by post-mortem analysis), possibly due to differences in AD models, brain region and/or plaque proximity. The J20 model has not been reported to exhibit significant levels of neurodegeneration; in line with this we did not observe any dystrophic neurites. Other studies have shown dystrophic neurites and dramatic synapse loss around plaques, however, these occurred within 15 µm of a plaque (Koffie et al., 2009). Here, neurites were imaged 50-200 µm from the nearest plaque edge, which would explain why no dystrophic neurites were observed. Therefore, the loss of axonal boutons was on cells, which did not degenerate during the experimental time frame.

This study presents novel *in vivo* measurements of axonal bouton density, revealing a significant reduction in TBs. This is consistent with a loss of synaptophysin staining found here (**Fig. 4h**) and in other similar studies (Spires et al., 2005). An explanation for the disparity in pre- and post-synaptic regulation could be that bouton loss occurs prior to spine loss in the cortex, where spines are subsequently upregulated to compensate for this, at least in the earlier stage of the disease. This ability may be lost in older animals and, thus, could explain the differences observed in the above-mentioned studies and our findings, with regards to spine density. Another explanation could be that axonal boutons are upregulated due to dendritic spine instability, but over compensation by the axons results in TB loss. Or, as a recent study has shown, Aβ is enriched only at the pre- and not at the post-synapse (Yu et al., 2018), which may also explain why, in our study, axonal boutons are more affected than dendritic spines. In addition, there could be underlying sex differences in these models of AD, which has not been extensively reported.

Further, the imbalance between the pre- and post-synaptic components, seen here, is consistent with what we have reported in the Tg4510 model of tauopathy (Jackson et al., 2017). Whilst this mismatch may be explained in tauopathy by the movement of pathological tau to the somatodendritic compartment (Hoover et al., 2010), the mechanisms of how Aβ causes synapse loss are less clear. Several studies have proposed candidate A receptors such as p75NTR (Knowles et al., 2009), Frizzled (Zagrebelsky et al., 2005) and LilrB2/PirB (Kim et al., 2013), all of which have been implicated in altered synaptic plasticity, but mainly on the post-synaptic side (Kim et al., 2013; McLeod et al., 2018; Zagrebelsky et al., 2005). Presumably, there is a soluble Aβ gradient, synapses close to amyloid plaques more likely to be lost through the toxic effects of soluble Aβ. Aside from this there could be a component of glial regulation implicated in this imbalance. Indeed, Hong *et al*. have shown that local activation of the innate immune system, via the microglial complement system, mediates early synaptic loss in AD, before apparent plaque formation, and importantly, inhibiting this response ameliorates synapse loss [13]. This further highlights the importance of glial involvement in AD-related synapse loss.

Another novel finding in this study was that, despite the stability of dendritic spine density, their survival was significantly reduced. On the other hand, spine turnover, driven by the formation of newly formed spines, was increased, presumably leading to the maintenance of stable spine density. Conversely, the significant reduction in TB density was paralleled by the highly significant reduction in survival fraction and loss of TBs, with only a modest, but significant, increase in gained TBs, which was not sufficient to maintain a stable TB density. The loss of presynaptic boutons but persistence of spine levels would be sufficient to account for the decline in input-output relationships observed in multiple neurophysiological studies of synaptic transmission in murine models of amyloidopathy (reviewed in (Randall et al., 2010)), although other explanations are feasible, for example, changes in action potential waveforms (Brown et al., 2011).

## Conclusions

In conclusion, our study reveals the unique, long-term growth dynamics of individual amyloid plaques and how this develops with disease progression. This is of particular importance when considering treatments that target amyoidopathy. In addition, our findings are consistent with previous work showing that synapses are targeted in amyloidopathy and we have shown, for the first time, that altered synapse dynamics leads to unstable synapses. Whilst studies are underway that seek to address the underlying pathology in AD and halt the disease, synaptic loss would still remain and continue to cause cognitive deficits. Thus, synaptic loss and instability would need to be addressed, alongside treating the underlying pathology, for cognitive benefits in AD.

## List of abbreviations

AD: Alzheimer’s Disease
APP: Amyloid Precursor Protein
FAD: Familial Alzheimer’s Disease
SSC: Somatosensory Cortex
RoR: Rate of rise
TBs: Terminaux boutons

## Materials and methods

### Animals

The J20 Aβ-overexpressing line carries a PDGF-β promoter-driven transgene for human APP, with *Indiana* (V717F) and *Swedish* (K670M – N671L) mutations. Both mutations are associated with FAD (Mullan et al., 1992; Murrell et al., 1991). Adult female mice of the J20 (TG; n = 15 animals) line and WT littermate controls (n = 15 animals) were used from 7 months (27 weeks) of age. Adult female mice from 11 months (45 weeks) (n = 14 animals) were also used for plaque analysis. This second set of animals was administered with a control vehicle for a drug discovery study, however, it was confirmed, with post-mortem immunohistochemistry, that this had no effect on measures of amyloid pathology (data not shown). All mice were given *ad libitum* access to food and water and maintained in a 12-hour light-dark cycle. All procedures were conducted by researchers holding a UK personal license and conducted in accordance with the UK Animals (Scientific Procedures) Act 1986 and subject to internal ethical review.

### Surgery

Cranial windows were surgically implanted over the SSC as previously described (Jackson et al., 2017). Briefly, mice were anaesthetized with isofluorane and administered dexamethasone (30 mg/kg) to limit brain swelling and the analgesic, buprenorphine (5 mg/kg), pre-operatively. The skull was exposed and a 5 mm diameter craniotomy drilled over the SSC. AAV serotype 2 expressing enhanced-GFP was stereotaxically injected into the SSC, ∼ 300 µm below the dura. A glass coverslip was placed over the craniotomy and sealed with glue and dental cement. A screw was placed in the skull on the contralateral side for added stability. The whole skull was subsequently covered in dental cement and a metal bar placed on top to allow head fixation at the two-photon microscope. Mice were allowed to recover for 3 weeks before imaging began. It was confirmed, by immunohistochemistry, that the cranial window did not cause an increase in amyloid pathology or microgliosis in any of the groups (data not shown).

### Imaging

5 mg/kg methoxy-XO4 (45% PBS, 45% propylene glycol, 10% DMSO) (2.9 mM) was injected intraperitoneally 24 hours prior to two-photon imaging to label the fibrillary beta sheet deposits of the amyloid plaques. 25 μl of dextran-Texas red was injected intravenously immediately prior to imaging to visualize blood vessels in some imaging sessions. A purpose built two-photon microscope equipped with a tunable coherent Ti:Sapphire laser (MaiTAI, SpectraPhysics) and PrairieView acquisition software was used for all imaging experiments. Mice were anaesthetized with isofluorane and secured to the microscope via the metal bar attached to the skull and a custom-built fixed support. Lacri-lube was applied to the eyes to prevent dehydration and temperature of > 34°C maintained by a heating blanket and rectal thermal probe. An Olympus 10X objective (NA = 0.3) was used to identify characteristic blood vessels to reliably relocate regions-of-interest (ROIs) at each imaging time point. In each animal an Olympus 40X (NA=0.8) water immersion objective was used to acquire several ROI stacks (300 µm x 300 µm, 512 x 512 pixels, z-step size = 3 µm for plaques or 75 µm x 75 µm, 512 x 512 pixels, z-step size = 0.5 µm for neurites). A pulsed 910 nm wavelength laser beam was used with a typical power at the sample of 35 mW.

### Analysis

Two-photon images were converted into stacks with ImageJ and the StackReg plugin run to align the GFP stacks in case of any movement. The GFP stacks were deconvolved with Huygens Deconvolution software using a quick maximum likelihood estimation with an experimentally-defined point-spread-function. The red channel stacks (with methoxy-X04 signal) were left unprocessed. Plaque images were denoised and maximum projections created. Plaque area was measured by manually outlining the total plaque and subsequently the core and measuring the area in ImageJ. The cloud area was calculated by subtracting the core area from the total plaque area. Prior to this all images were blinded using a custom ImageJ plugin so that time points were unknown at the point of analysis. The average rate of rise (RoR; µm^2^/week) of each plaque, was measured as the slope of a linear function interpolated for the plaque-to-plaque surface value in time (weeks). For dendritic spine and axonal bouton analysis each time-point was registered to the second imaging session using MIPAV and converted to a 4D stack using a custom ImageJ macro. Each neurite to be analyzed was traced and the length measured using another custom macro. Subsequently, the spines or boutons were manually counted using the cell counter ImageJ plugin. Spines/boutons were counted in 3D image stacks. while maximum projections are shown in the figures. Only TBs, not en passant boutons, were counted and identified as protrusions emanating from side of the axon with a head and neck. From this the density (μm^−1^) and TOR (survival, as a fraction of the first time-point) were calculated. Statistics were performed using GraphPad Prism 7 or Sigma Plot 13. RM ANOVA comparisons (with Holm-Sidak method post-test) were utilized where multiple comparisons were made, unless otherwise stated.

### Research data

Due to confidentiality agreements with research collaborators, supporting data can only be made available to bona fide researchers subject to a non-disclosure agreement. Details of the data and how to request access are available on request to the corresponding author

## CLARITY Staining

Transcardiac perfusion with 4% paraformaldehyde was performed in a separate cohort of J20 animals at 10 months of age. Brains were sliced in 1 mm sections (Brain matrix, Alto) after 1 to 5 days incubation in 4% paraformaldehyde. To preserve molecular information and structural integrity brain slices were incubated with a hydrogel solution (4% acrylamide and 0.25% VA-044 in 1X PBS). Therefore, the tubes were degassed and the hydrogel was allowed to polymerise for 3 h at 37°C. The slices were transferred into clearing solution (SDS, sodium borate) at 37°C using the X-CLARITY system (Logos Biosystems). The slices were placed in a tube containing 1 ml of 5% BSA. The required amount of primary antibody was pipetted directly in the BSA solution and incubated for 4 days at 37°C. Excess of antibody was washed away with PBS for 3 x 2 h at 37°C. Secondary antibodies diluted in 5% BSA were incubated for 4 days at 37°C and excess removed by washing the slices with PBS for 3 x 2 h at 37°C. Concentric circles 5 µm apart were drawn on maximum projection images around the plaque and the mean fluorescence measured radiating out from the edge of the plaque.

---

**Supplementary Figure 1.**
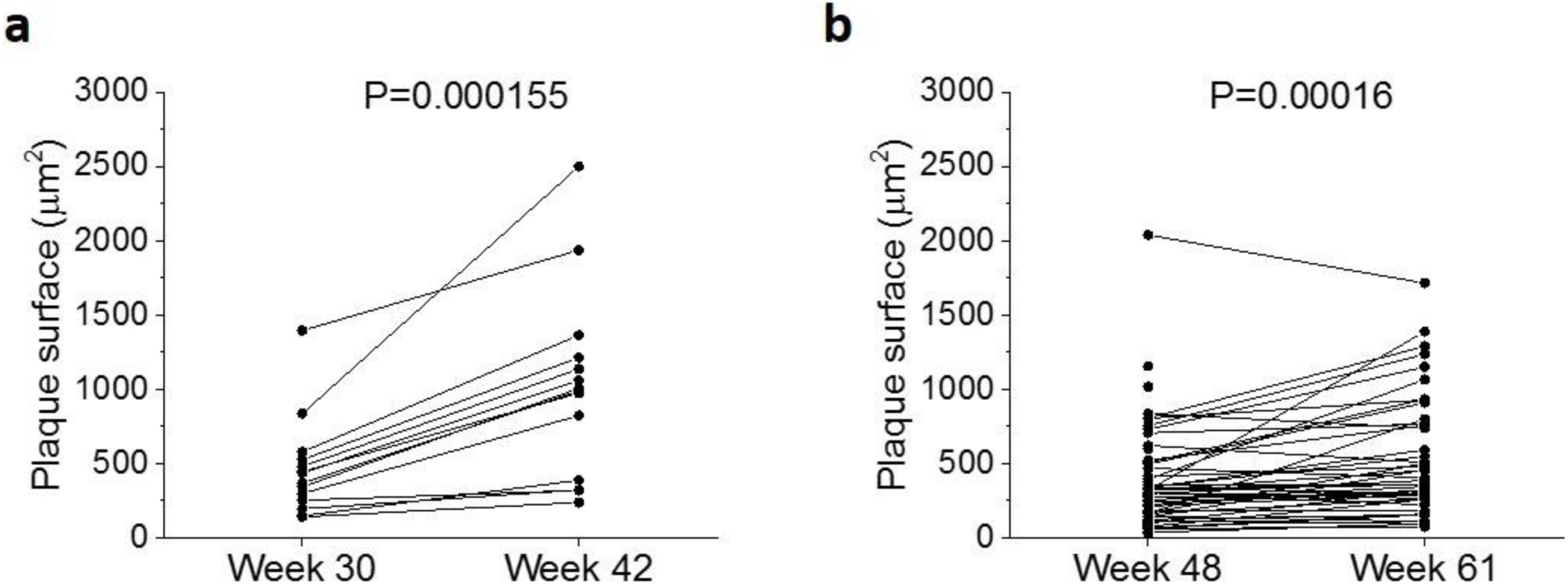
Surface area of individual amyloid plaques in J20 mice in the early (a) and late (b) groups. P values calculated using paired t-tests (these values have not been corrected for multiple comparisons as they are 2 orders of magnitude below the threshold of 0.05).

**SI Table 1.**
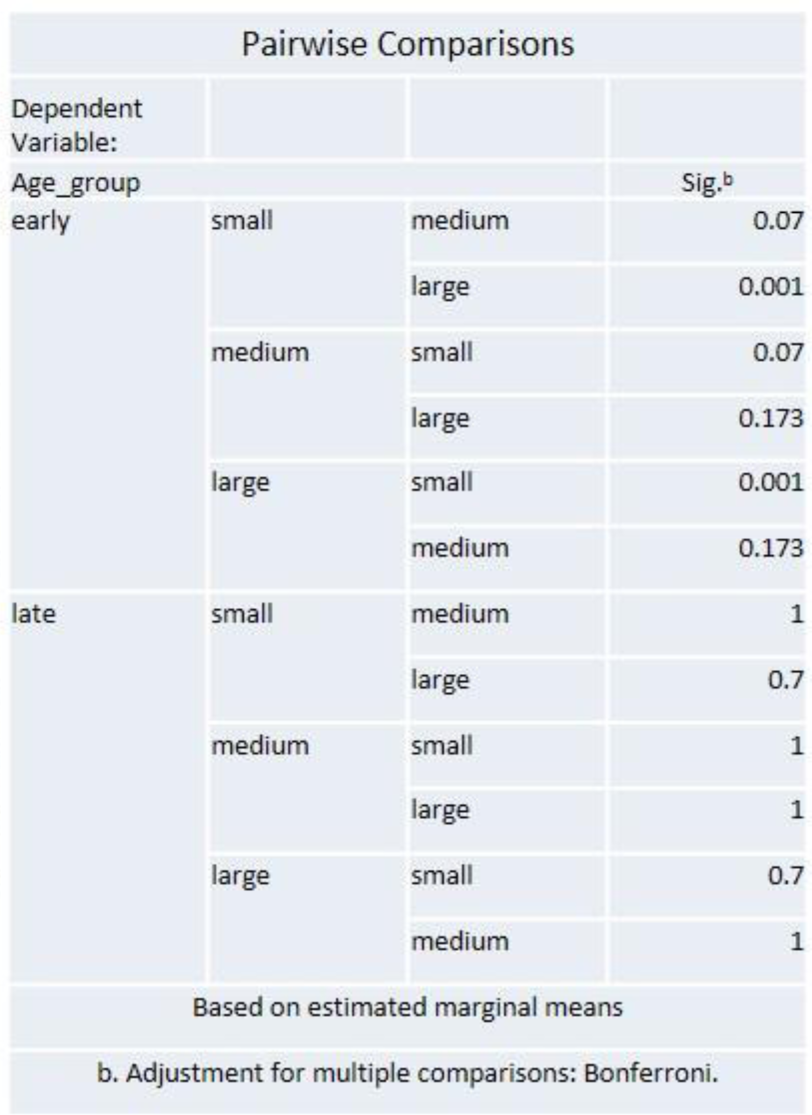
Pairwise comparisons between age-groups and plaque-initial-size groups. Top. In younger animals, small plaques grew significantly faster, in comparison to both medium and large ones. No difference was however observed between medium and large-sized plaques. Bottom. No effect of plaque’s initial size on growth rate was observed at a later age-stage.

| Pairwise Comparisons |  |  |  |
| --- | --- | --- | --- |
| Dependent Variable: |  |  |  |
| Age_group |  |  | Sig.<sup>b</sup> |
| early | small | medium | 0.07 |
|  |  | large | 0.001 |
|  | medium | small | 0.07 |
|  |  | large | 0.173 |
|  | large | small | 0.001 |
|  |  | medium | 0.173 |
| late | small | medium | 1 |
|  |  | large | 0.7 |
|  | medium | small | 1 |
|  |  | large | 1 |
|  | large | small | 0.7 |
|  |  | medium | 1 |
| Based on estimated marginal means |  |  |  |
| b. Adjustment for multiple comparisons: Bonferroni. |  |  |  |

**SI Table 2.**
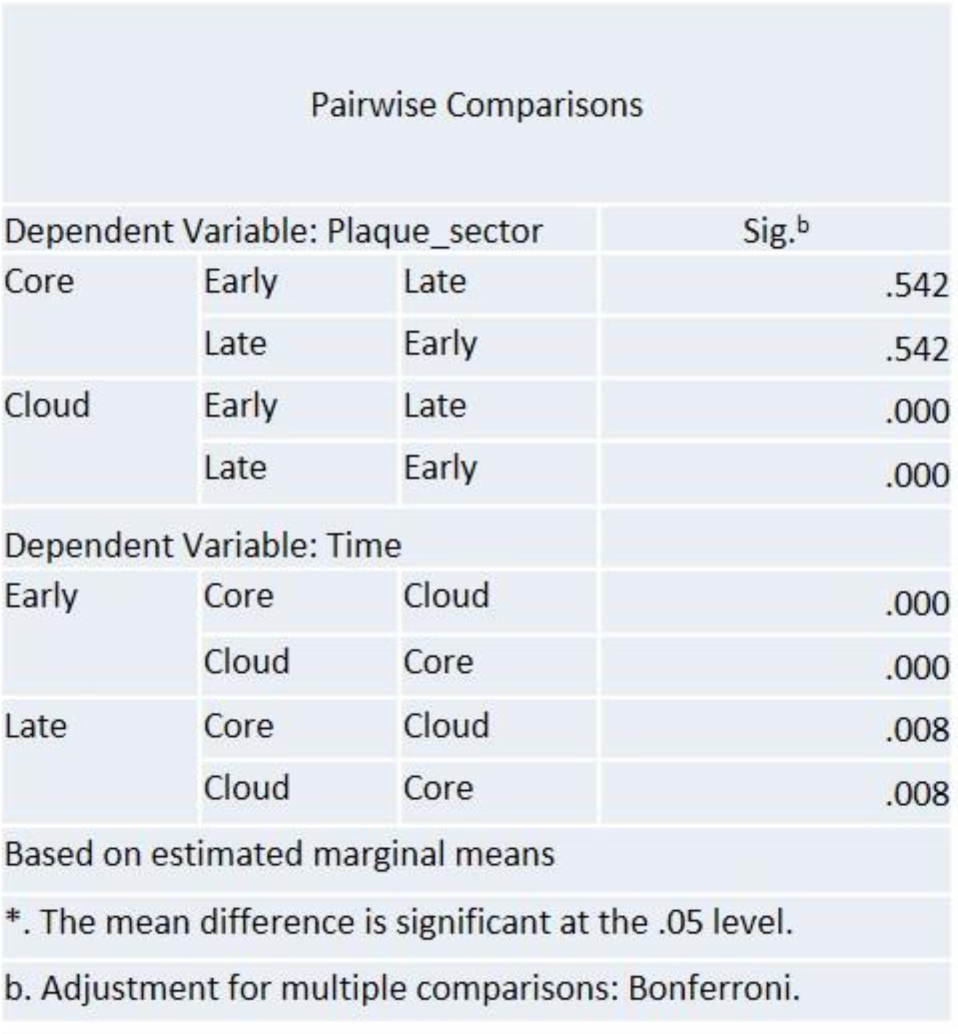
Pairwise comparisons between age-groups and plaque-sectors. Top. The core of the plaque does not grow at different rates between younger and older animals; however, the cloud sector grows faster in younger animals. Bottom. Both in younger and older animals, the cloud sector grows faster than the core.

| Pairwise Comparisons |  |  |  |
| --- | --- | --- | --- |
| Dependent Variable: Plaque_sector |  |  | Sig.<sup>b</sup> |
| Core | Early | Late | .542 |
|  | Late | Early | .542 |
| Cloud | Early | Late | .000 |
|  | Late | Early | .000 |
| Dependent Variable: Time |  |  |  |
| Early | Core | Cloud | .000 |
|  | Cloud | Core | .000 |
| Late | Core | Cloud | .008 |
|  | Cloud | Core | .008 |
| Based on estimated marginal means |  |  |  |
| *. The mean difference is significant at the .05 level. |  |  |  |
| b. Adjustment for multiple comparisons: Bonferroni. |  |  |  |

## Declarations

### Authors’ contributions –

JSJ, FT, AR, MLH and MJON designed the study, TS analysed the in vivo data; FT and JSJ carried out the in vivo experiments; JP carried out the CLARITY; KS, JH, AO-E, TKM and ZA characterized the model; AR, MJON, JSJ jointly supervised the project.

